# Exploring multi-omics latent embedding spaces for characterizing tumor heterogeneity and tumoral fitness effects

**DOI:** 10.1101/2023.07.05.547886

**Authors:** Fengao Wang, Junwei Liu, Feng Gao, Yixue Li

## Abstract

The ecological and evolutionary perspectives of tumorigenesis can be characterized as a process of microevolution in tumor cells that altered the tumor microenvironment and further induced tumor cell proliferation, metastasis, and the death of tumor patients. Here, we introduced XgeneVAE, an interpretable unsupervised deep learning framework that quantified the semantic changes in multi-omics embedding space for characterizing the microevolution processes and fitness effects of heterogeneous tumor samples. We then validated that the scales of the latent embedding variances can reflect the differences in the overall survival of tumor patients, as well as their applications in uncovering the driving genomic alternations in different cancer types. These results confirmed that the XgeneVAE model can better represent the heterogeneity in distinct cancer types and as an interpretable model for understanding the fitness effects in tumorigenesis and their association with clinical outcomes.

## Introduction

Tumorigenesis can be characterized as a aggregations of changes that have occurred in the DNA sequence of the genomes of cancer cells and further effects its proliferation and differentiation [1]. Recently, this process has increasingly been accepted as an evolutionary process and populations of cancerous cells evolve, much like populations of organisms do. This microevolutionary process involves genomic alternations in tumor cells that, under natural selection, can induce changes in both the tumor microenvironment and phenotype [2]. These alterations are captured and reflected via multi-omics data at the molecular level, which includes genomics, epigenomics, transcriptomics, and proteomics. Each of these type of datasets provides unique insights into the tumoral cellular microenvironment [3]. Given the large-scale availability of this multi-omics data, a primary challenge in the field is how to effectively integrate these data and the extract valuable features from such rich resources [4].

Recently, deep learning has emerged as a powerful approach for extracting non-linear informative features from multi-omics data in oncology [5-7]. However, the majority of these studies have relied on supervised learning methods, given the deep learning’s inherent design for learning from experience and mapping input data to the distribution of supervisory labels [8]. Furthermore, a significant drawback in most of these studies is the lack of consideration for the complexity of tumor microevolution, which results in poor biological interpretability [9, 10].

Recent advancements in unsupervised methods have demonstrated that pretraining procedures can extract universal features and unveil novel knowledge from training data [11, 12]. Previous studies have successfully utilized unsupervised methods such as gene embedding and variational autoencoders (VAEs) to extract latent factors from gene expression or multi-omics data [13]. Such latent factors have been shown to be biologically relevant [8, 14, 15]. Unsupervised autoencoder models can encode input data into a semantic embedding space. Following the microevolutionary biology of tumors, the distance in the embedding space can signify the variance or evolutionary distance and change of tumor microenvironment in the process. This concept of an embedding semantic space originates from natural language processing, where sentences are mapped to a continuous vector space based on their distributional properties. This approach has produced impressive results, preserving the semantic relationships between words and phrases as distances within the embedding space. In this study, we hypothesize that multi-omics features also follow a distributional pattern. The characteristics of multi-omics data can be defined by their associations in terms of multi-omics features, and the embedding space can be used to encapsulate the entirety of cancer’s complexity [16]. In this space, each patient is represented by a point, with semantically similar patients situated closely to each other [17].

In this study, we introduce an unsupervised pretraining framework for multi-omics data, XgeneVAE, along with a semantic change metric of multi-omics embedding space from the microevolutionary process perspective. This metric is used to characterize the intratumor heterogeneity and predict the phenotype of cancer patients. Our Variational Autoencoder (VAE)-based framework can reduce the dimensionality of high-dimensional multi-omics data and encode it to represent the embedding features of tumors [18]. We computed the semantic changes of tumors and identified gene mutations that significantly contribute to these changes. Drawing inspiration from the Phenotype Embedding (PE) theorem (Liu, et al., 2023), we used the semantic embedding change to predict the phenotype of patients, and applied overall survival as the ground truth reference. Remarkably, our methods successfully predicted the overall survival of cancer patients (log-rank test P-value<0.05) without any clinical information used in the training. The results show the semantic change in the multi-omics embedding space is related to biological phenotype and cancer heterogeneity. Our study introduces a novel methodology and provides fresh insight into the heterogeneity and fitness of tumors by computing the semantic change in the embedding space.

## Results

### Unsupervised pretrain model to learn the semantic embedding

In this study, we introduce XgeneVAE, an unsupervised pretraining framework for multi-omics data, accompanied by a unique metric methodology for computing semantic changes within the embedding space to elucidate the heterogeneity and phenotype of cancers, as shown in Figure 1 A. XgeneVAE was pretrained on the TCGA pan-cancer dataset, and this process did not incorporate any supervised labels [19]. A key step in pretraining involved reducing the dimensionality of multi-omics features to obtain a representative semantic embedding of the data, enabling the model to encapsulate the inherent structure and relationships within the multi-omics data [20].

**Figure 1.**
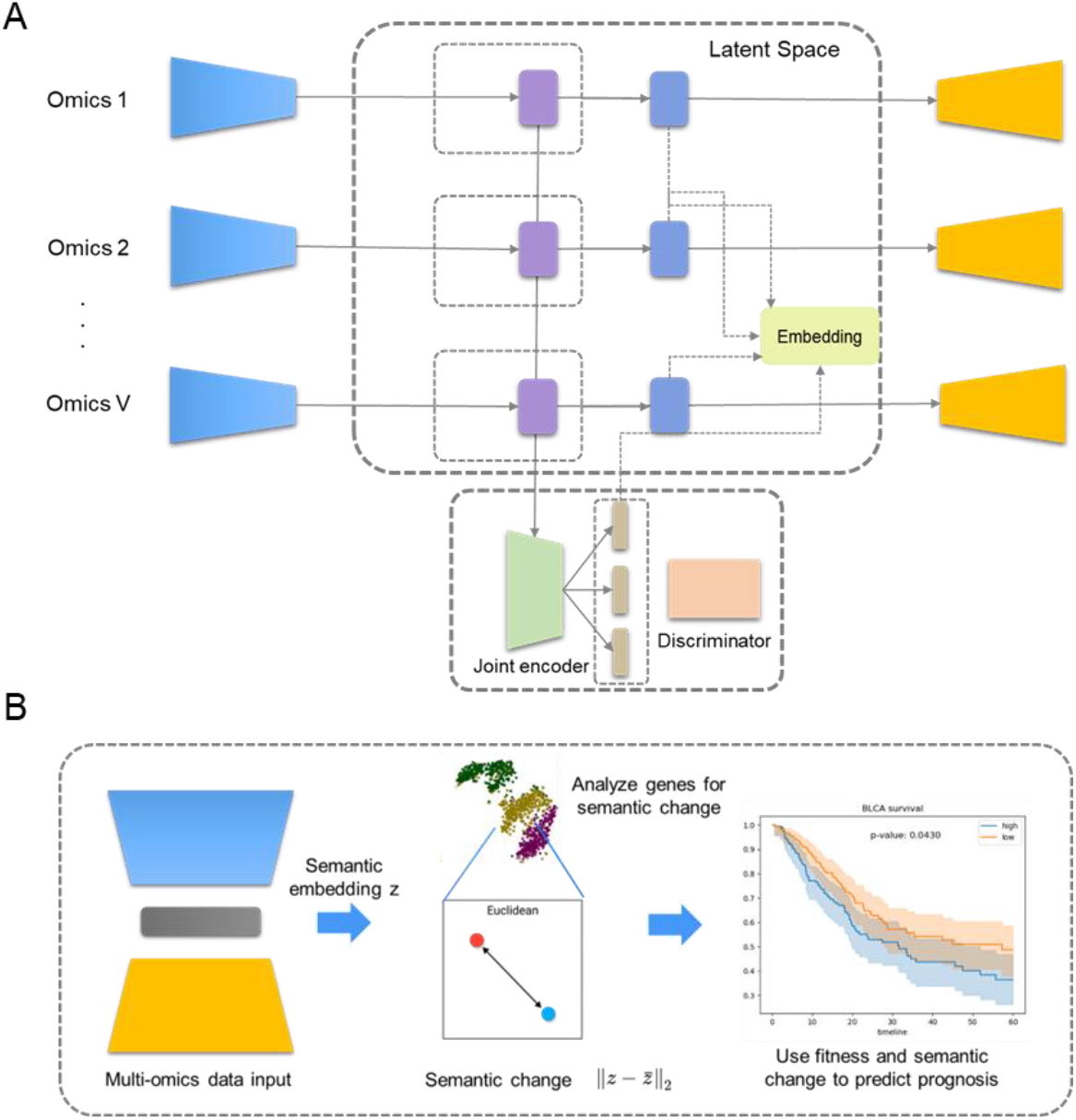
Framework of XgeneVAE(A) and semantic change in multi-omics embedding space(B)

We scrutinize this low-dimensional embedding latent space and visualize tumor clustering using t-SNE for the direct reduction of embedding latent variables to a two-dimensional format. The multi-omics features of 8,174 samples, as learned by XgeneVAE, are presented in a scatter plot in Figure 2. The visualization results showcase a hierarchical structure and a clear demarcation between different cancer types. Intriguingly, cancers originating from the same system or organ are spatially close yet distinguishable in the visualizations, as depicted in Figure 2. This indicates that these cancer points in the embedding space are semantically similar, underscoring the effectiveness of our approach.

**Figure 2.**
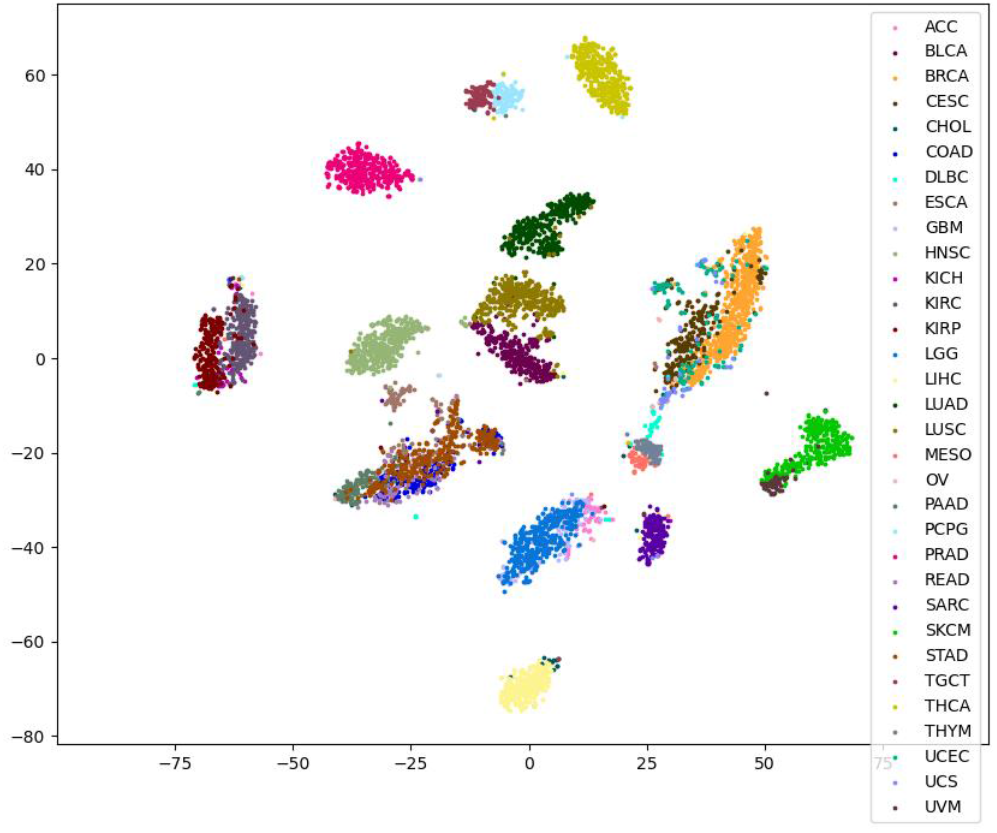
Visualization of tumor clustering by using t-SNE for the direct reduction of multi-omics embedding latent variables to a two-dimension. Samples of various tumor types were plotted with different colors shown in the figure.

In the clustering visualization, three subtypes of kidney tumors - kidney clear cell carcinoma (KIRC), kidney papillary cell carcinoma (KIRP), and kidney chromophobe (KICH) - are clearly differentiated. Yet, they cluster closely together due to their shared organ of origin. Similarly, despite being distinct in the visualization, two subtypes of lung cancer cluster closely on a larger scale. Cancers from the digestive system, although separated based on their subtypes, also cluster near each other in the broader landscape. This spatial distribution mirrors the shared and unique molecular characteristics of cancers based on their tissue or organ of origin.

To validate the feasibility of the learned embedding relationship between samples, we used cancer types as a readily available ground truth reference to examine the embedding features. We applied Principal Component Analysis (PCA), t-Distributed Stochastic Neighbor Embedding (t-SNE), and Uniform Manifold Approximation and Projection (UMAP) for comparison with our model’s embedding [21-23]. While both t-SNE and UMAP are capable of capturing non-linear relationships, PCA is a linear method for dimension reduction. These methods were used to embed the same multi-omics data into a two-dimensional space.

The low-dimensional representations learned by PCA were plotted on a scatter graph, as shown on the right of Figure 2. The 2D embedding obtained from PCA displays a more mixed clustering structure compared to that learned by XgeneVAE, making it challenging to discriminate between different tumor types. We then input this 2D representation into XGBoost to classify 32 classes of tumors [24].

We measured the classification performance of the four methods using four multi-class classification metrics: overall accuracy, weighted precision, weighted recall, and weighted F1 score. The performance results of 5-fold cross-validation are displayed in Table 1. As shown, the 2D representation of embedding learned by XgeneVAE outperforms those 2D dimensions learned by the other methods. This suggests that the embedding space learned by XgeneVAE effectively represents the multi-omics data and accurately reflects the heterogeneity between cancers.

**Table 1.**
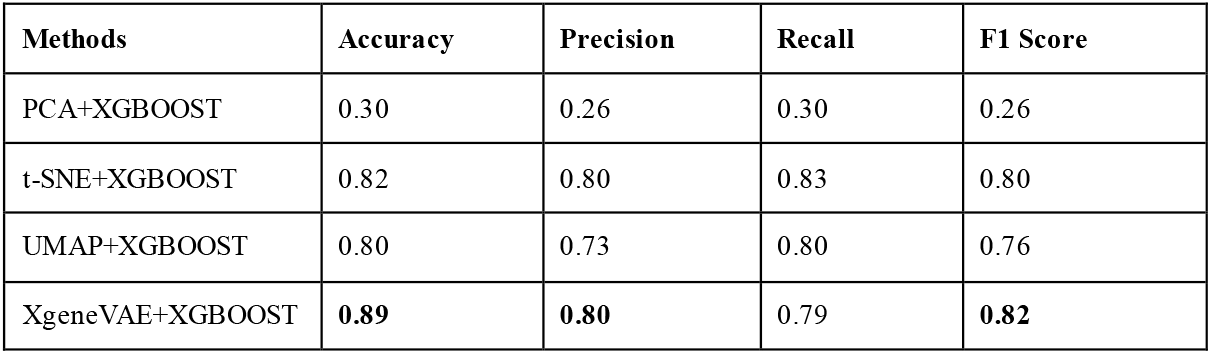
Cancer classification performance of multi-omics embedding learned by different dimensionality reduction methods and XgeneVAE.

### Learn the tumor heterogeneity of multi-omics data in the embedding latent space

Based on the embedding space of multi-omics data, and drawing inspiration from the Phenotype Embedding (PE) theorem [25] and as well as the semantic meanings within natural language processing, the values derived within the embedding space can be utilized to represent macroscopic phenotypes. Here, we consider the euclidean distance between two points as a semantic change in the embedding space:

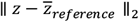

as a metric for quantifying the heterogeneity and fitness effects of tumor multi-omics data, as shown in Figure 1 B. This is defined by the multi-omics embedding of a tumor sample within the embedding space relative to a reference point. This reference point could be any randomly selected point within the embedding space, where each point embedding reference a unique biological insight. The euclidean distance to the reference point symbolizes the deviation within the embedding space, which is determined by the input multi-omics data. The map between the input data and the embedding features was learned during the pretraining stage.

### Semantic change driven gene mutation identification of intratumor heterogeneity

Intuitively, our objective is to identify mutations that induce significant semantic changes relative to the reference point within the embedding space. Initially, to focus on the internal structure of cancer, we employ the k-nearest neighbors (KNN) algorithm to compute the average centroid for each type of cancer sample within the embedding space [26]. This KNN-derived centroid effectively represents the mean features of a given cancer type within the embedding space. We select the embedding centroid of each type of cancer sample as the reference point. For each individual cancer sample embedding z, we calculate the euclidean distance from each sample to this average point within the embedding space:

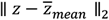

For each patient, this semantic deviation in the embedding space reflects the intratumor heterogeneity. When considering a group of patients, the mean euclidean distance represents the evolutionary difference within the cancer type. Subsequently, we apply Integrated Gradients (IG), a gradient-based attribution method, to compute the contribution score of each feature within the multi-omics data to the semantic change [27]. This process generates a score quantifying the influence of each feature on the semantic shift within the embedding space. This score offers an interpretable metric for delineating how individual molecular multi-omics features impact the heterogeneity and conservation of cancers. As shown in Figure 3, we enumerate the top 10 gene mutations most relevant to the corresponding cancer semantic changes, thereby demonstrating the interpretability of XgeneVAE and semantic shifts by revealing the contribution scores for each gene mutation.

**Figure 3.**
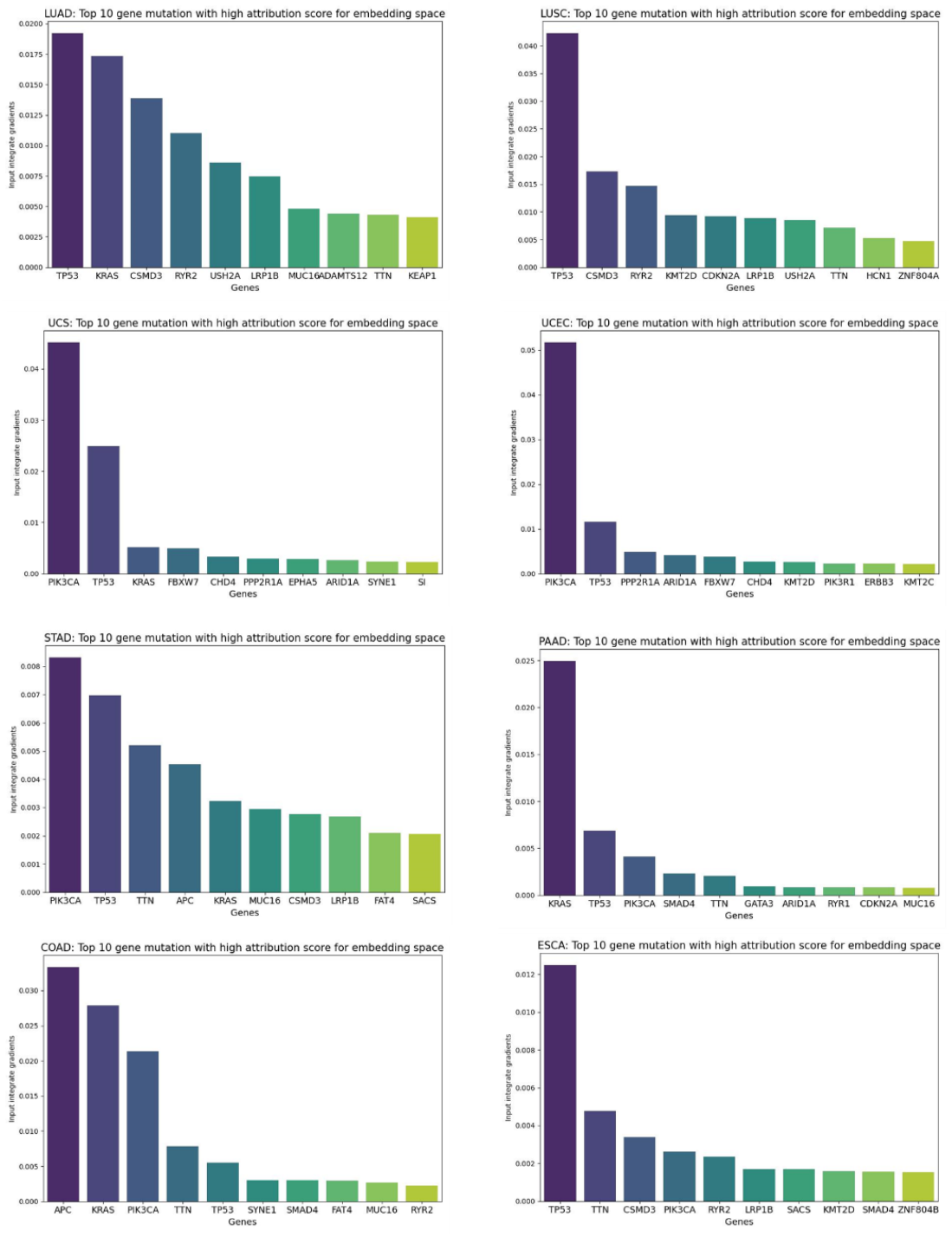
Top 10 gene mutations significantly contributing to the semantic changes observed in a particular cancer type. Using Integrated Gradients (IG), we calculated the contribution scores of each feature from multi-omics data to the observed semantic changes per sample. IG values were subsequently computed for numerous samples and averaged to ascertain the average feature importance associated with each gene.

Most of the genes we identified are widely associated with cancer development and are frequently corroborated in existing literature. To validate the top genes revealed by XgeneVAE, we first compared the genes, ranked by contribution for each tumor type, with genes associated with the corresponding tumor type from GeneCard [28]. We chose GeneCard because of its comprehensive disease gene sets, integrated from roughly 150 different web sources, thereby covering the majority of tumor types in our analysis. We compared the genes at 100 different thresholds, spaced evenly from 1 (the most important gene) to 58,043 (the total number of genes). A total of 21 tumor types had gene sets found in GeneCard and were thus chosen for analysis. The overlap of GeneCard demonstrated that tumor driven genes identified by XgeneVAE are biologically relevant.

### Predict the phenotype of overall survival by semantic change

Beyond the intrinsic differences within the embedding space of cancer, we also consider the distance from each feature point to the mean of normal tissues. This distance reflects the deviation between tumor cells and normal cells at the molecular level, a deviation caused by multi-omics and changes in the microenvironment. We employ the KNN algorithm to compute the mean of the normal samples embedding feature. This provides a representative average of the embedding characteristics within normal tissues. For each feature point, the semantic change or deviation in the embedding space relative to the reference can be calculated euclidean distance for each sample:

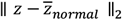

This deviation from normal tissue is considered to reflect the level of severity and fitness of the tumor in the evolutional process. In this context, the fitness value is inversely correlated with the patient’s survival phenotype, with a higher semantic change indicating a higher risk score.

To examine the relation semantic change with tumor fitness and patient survival, the overall survival event and time of patients were used to validate as the ground truth reference. We calculated the semantic change value for each patient’s embedding point, relative to the average point for normal tissue. We identified the median semantic change value within each cancer group and utilized it to categorize samples into high-risk and low-risk groups. Subsequently, we visualized the survival outlook for cancer patients through Kaplan-Meier (KM) survival curves, as depicted in Figure 4. Statistical validation results substantiate that the semantic changes in the embedding space can effectively differentiate patients in the majority of cancers (log-rank test P-value<0.05). This suggests that the semantic change relative to the average point for normal tissue in the embedding space can be used to predict fitness of tumors and the prognosis of patients.

**Figure 4.**
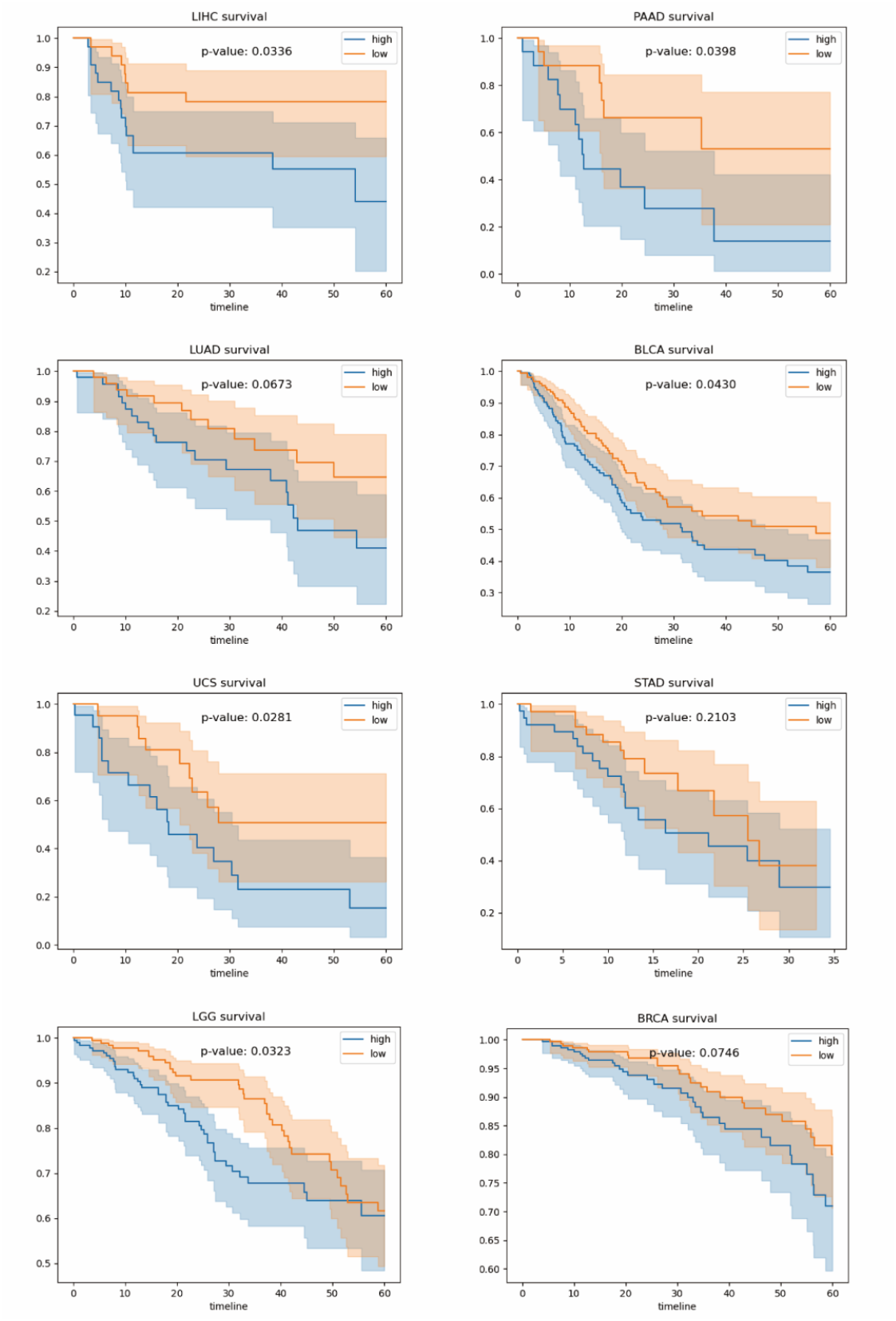
Kaplan-Meier (KM) survival curves of low- and high-risk patients via semantic changes learned by XgeneVAE. Low and high risks are defined by the median 50% percentile of semantic change compared to the mean embedding point of normal tissues. Log rank test was used to test for statistical significance in survival distributions between low- and high-risk patients.

## Conclusion

In this study, we introduced XgeneVAE, an unsupervised pretrain framework, along with a novel semantic change metric of embedding space to characterize cancer and predict the phenotype of cancer patients. Drawing inspiration from the Phenotype Embedding (PE) theorem [25], this approach serves as a new metric to characterize the heterogeneity of cancers at the level of multi-omics data and to compute the fitness effects of multi-omics. To our knowledge, this is the first use of semantic change in the embedding space for identifying cancer-driving genes and predicting phenotype in an unsupervised manner from the perspective of the tumor microevolutionary process. We trained XgeneVAE on TCGA pan-cancer datasets and obtained a representation of multi-omics data within the embedding space. The results show the semantic change in the multi-omics embedding space is related to biological phenotype and cancer heterogeneity. Our approach demonstrates substantial potential for uncovering novel biomedical insights from unsupervised deep learning models, with findings that align well with current domain knowledge, including the biological annotation and academic literature.

## Materials and Methods

### Datasets collection and preprocessing

We sourced multi-omics data for 32 pan-cancers from the TCGA database, which consisted of 8,174 samples, including normal tissue samples. Each cancer dataset incorporated four types of omics data: RNA-Seq gene expression profiles, DNA methylation profiles, gene mutation data, and copy number variation (CNV) data. Additionally, corresponding clinical information was included.

For multi-omics data, in line with previous studies [29], we included five types of nonsynonymous mutations: missense and nonsense mutations, frameshift insertions and deletions, and splice sites. Within the TCGA dataset, only high-confidence mutations that passed the Multi-Center Mutation Calling in Multiple Cancers filter were considered. We included 4,539 genes that were mutated in at least 1% of TCGA samples. Gene expression levels were represented by log_2_(TPM + 1), where TPM denotes the number of transcripts per million, using data from 9,806 TCGA tumors. We included 6,016 genes that showed a mean and standard deviation greater than 1 in the TCGA samples. For DNA methylation, data from 12,039 TCGA samples were used. We excluded probes with low methylation (β value, <0.3) in more than 90% of TCGA samples, retaining 6,617 probes for further analysis. CNA data, collected from Affymetrix SNP 6.0 arrays of 11,101 TCGA samples, were mapped onto approximately 310,000 consecutive segments of length 10,000 bases along the chromosomes, with each CNA being weighted by the percentage covered by the CNA. We included 7,460 informative CNA segments, which fulfilled the criteria of zeros in less than 5% of samples, a mean of absolute values greater than 0.20, and a coefficient of variation greater than 0.20 in TCGA. In our study, mutation and expression data were gene-based, while methylation and CNA data were represented by probes and chromosomal segments, respectively.

### Multi-omics VAE

The overall structure of our model is depicted in Figure 1. A Variational Autoencoder (VAE) is a potent deep generative model capable of learning essential information from the manifold of multi-omics data [30]. Building upon the fundamental VAE, we proposed a framework for learning embeddings from multi-omics data.

In this framework, we deploy omic-specific variational autoencoders to learn low-dimensional omics-specific tumor embeddings. Concurrently, we employ a shared feature encoder to learn the joint tumor embedding [31]. The embeddings from all omics layers are then aligned and fused to derive a multi-omics embedding, which serves as the overall tumor embedding. To achieve alignment across omics-specific data spaces, we use a discriminator (D). This aligns the tumor embeddings across different omics layers using adversarial learning, essentially harmonizing the multi-omics data representations. This alignment is critical to ensure that our VAE can appropriately analyze and interpret the multi-omics data, leading to more meaningful and accurate biological insights. Then we concatenate all the omics specific feature and the shared feature as the embedding of multi-omics data in the late fusion method [32].

### VAE Framework

Following the based VAE, the VAE network was optimized by maximizing the variational evidence lower bound (ELBO) defined in Equation:

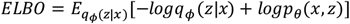

Equation can further transform to Equation 3:

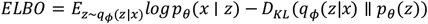

The network was optimized to maximize Equation 3 and for the deep learning model, the generative distributions can be implemented by the deep decoder neural network, and the variational posteriors by deep encoder neural networks. The VAE loss can be defined as follows:

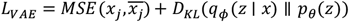

For multi-omics data, each omics data was first passed into VAE encoder to reduce the dimensionality from high dimensionality to 64 in the embedding space. For the omics input **x**, the VAE encoder network outputs two vector, the mean vector *μ* and the standard deviation vector *δ*, which determined the Gaussian distribution *N*(*μ, δ*) of the latent variable **z** in the embedding space. The latent variable z is randomly sampled from the distribution. In order to achieve backpropagation in deep neural network, the reparameterization trick was applied using Equation (1) to approximate the latent variable z:

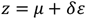

Then the latent variable **z** was passed to reconstruct the omics data 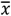 of the input *x* through the VAE decoder network.

### Cross modal adversarial learning

Apart from the fundamental Evidence Lower Bound (ELBO) computations, we’ve introduced a cross-modal adversarial learning procedure to align the omics-specific representations within the embedding space. In this framework, shared encoders are used to extract common features from various omics data. These common features are then processed by a discriminator that discerns the source omics data for these features. Through this process, we aim to extract and emphasize the shared, joint features from diverse omics data.

In the latent space, the omics-specific embeddings are again fed into the discriminator, which attempts to identify the omics data inferred from these embeddings. Consequently, the encoders are trained adversarial to deceive the modality discriminator. This adversarial training procedure ensures the alignment of embeddings from different omics data, promoting a more comprehensive and unified understanding of the multi-omics landscape [33]. By aligning and integrating these various omics-specific representations, our model facilitates a more coherent and informative multi-omics analysis:

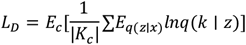

### Visualization of embedding dimension

Embedding matrices were visualized by either t-distributed stochastic neighbor embedding (t-SNE) or PCA. Three-dimensional t-SNE was implemented with perplexity = 5 and 50,000 iterations. Three components PCA were implemented using default setting. Both t-SNE and PCA were done using scikit-learn library.

### Interpretability of Multi-omics Data

By leveraging Integrated Gradients (IG), a gradient-based attribution method, we are able to assign a contribution score to each feature present in our multi-omics data, including individual gene mutations. These scores essentially quantify the influence that each feature has on the semantic shift observed within the embedding space. In doing so, we’re provided with an interpretable metric that clearly delineates how each feature, be it a gene mutation or another form of multi-omics data, contributes to the heterogeneity and conservation observed within cancers.

### Implementation

We adopted a 5-fold cross-validation strategy to partition all datasets, ensuring that each split was stratified according to the sample collection site to help control potential batch artifacts. Our model, XgeneVAE, was developed on the PyTorch framework, with the interpretability analysis being performed using Captum. We employed the Adam optimizer to improve XgeneVAE, with a learning rate of 0.0001, a batch size of 64, and 0.001 weight decay. The training process was finished on the NVIDIA RTX 3090 with CUDA acceleration. The validation set was used to select the model with the best performance during the pretraining stage. The effectiveness of the pretraining stage was assessed using the LogME method, while the semantic change analysis was conducted in an unsupervised manner on the training dataset.

